# Neurogenesis-mediated forgetting of complex paired-associates memories

**DOI:** 10.1101/2021.03.25.437009

**Authors:** Jonathan R Epp, Leigh C.P. Botly, Sheena A Josselyn, Paul W Frankland

## Abstract

The hippocampus is a critical structure involved in many forms of learning and memory. It is also one of the only regions in the adult mammalian brain that continues to generate new neurons throughout adulthood. This process of adult neurogenesis may increase the plasticity of the hippocampus which could be beneficial for learning but has also been demonstrated to decrease the stability of previously acquired memories. Here we test whether increased production of new neurons following the formation of a gradually acquired paired-associates task will result in forgetting of this type of memory. We trained mice in a touchscreen-based object/location task and then increased neurogenesis using voluntary exercise. Our results indicate that mice with increased neurogenesis show poor recall of the previously established memory. When subsequently exposed to a reversal task we also show that mice with increased neurogenesis require fewer correction trials to acquire the new task contingencies. This suggests that prior forgetting reduces perseveration on the now outdated memory. Together our results add to a growing body of literature which indicates the important role of adult neurogenesis in destabilizing previously acquired memories to allow for flexible encoding of new memories.

## Introduction

In the subgranular zone of the hippocampus, new neurons are continuously generated throughout adulthood. Over the course of weeks, these new neurons develop into mature and functional granule cells. As early as 1-2 weeks following proliferation, new neurons begin to synaptically integrate into the existing hippocampal circuitry(Markakis and Gage, 1999; Toni et al., 2007, 2008). Once incorporated, they are thought to contribute to ongoing hippocampal function including learning and memory, particularly in the spatial and contextual domains. Adult generated neurons mature and eventually become indistinguishable from developmentally generated granule cells(van Praag et al., 2002; Espósito et al., 2005; Laplagne et al., 2006; Zhao et al., 2006; Stone et al., 2011). A number of studies have suggested that enhancing hippocampal neurogenesis facilitates the encoding of hippocampal memories. Interestingly, nearly all of these studies have utilized a similar experimental design in which neurogenesis is manipulated before behavioural training occurs. Voluntary exercise, pharmacological manipulations as well as transgenic mouse models with elevated neurogenesis have all been associated with some form of cognitive enhancement(Creer et al., 2010; Sahay et al., 2011; Bolz et al., 2015). Conversely, it has been shown that decreasing neurogenesis impairs hippocampal memory encoding. However, there are also exceptions to these studies in which increased neurogenesis is not linked to improved learning and memory suggesting that only some tasks or components of memory benefit from increased neurogenesis(Clark et al., 2008; Kohman et al., 2012; Swan et al., 2014).

These experiments, which involve manipulation of neurogenesis prior to behavioural examination, typically posit that adult generated neurons confer additional properties to the hippocampus that are otherwise absent. At the cellular level this has included ideas such as enhanced excitability while at the behavioral level enhanced discrimination ability. Such an explanation, while interesting, overlooks the fact that the continuous integration of new neurons must necessarily remodel the existing hippocampal circuits. We have predicted and demonstrated that this integration is disruptive to existing memories that are represented, at least in part by the existing connectivity of the hippocampus(Frankland et al., 2013; Akers et al., 2014; Epp et al., 2016; Ishikawa et al., 2016; Gao et al., 2018; Cuartero et al., 2019; Scott et al., 2021). At face value, this prediction is somewhat counterintuitive given the wealth of literature suggesting that neurogenesis is beneficial to hippocampal memory encoding. However, it is not incompatible these findings. We have previously demonstrated that pre-training manipulation of neurogenesis (enhancement using voluntary running), enhances the rate of acquisition of a spatial version of the Morris water task but had no impact on the expression of the spatial memory during a probe trial conducted following training. However, if spatial training precedes the increase in neurogenesis, we show that later memory retention is impaired(Epp et al., 2016). Importantly, we have also demonstrated that the same mice that show impaired retention due to enhanced neurogenesis also exhibit facilitated reversal learning compared to mice with normal levels of neurogenesis. This points to an important functional consequence of adult neurogenesis. That is, the incorporation of new neurons acts to reduce interference between old and new memories by weakening the old memories. Therefore, neurogenesis exerts a pro-cognitive effect because one of its primary actions causes forgetting.

Here, to extend our understanding of the range of memory types that are impacted by post-training manipulation of neurogenesis, we examined the impact of voluntary running on the retention of a complex paired associates learning (PAL) task acquired in a touch screen operant conditioning platform (Roebuck et al., 2018). We demonstrate that voluntary running, increases neurogenesis and weakens the retention of previously acquired memories, even those acquired gradually over a long period of training. Further, we provide supporting evidence that this type of forgetting may be beneficial for new conflictual learning.

## Materials and Methods

### Mice

We used wild-type mice that were derived from an f1 cross between C57Bl/6N and 129Svev mice (Taconic). Breeding was conducted in the Hospital for Sick Children animal facility. Mice were housed in standard laboratory conditions with 4 mice per cage. The housing rooms were maintained on a 12 h light/dark cycle and behavioural testing was conducted during the light phase. All experiments were approved by the animal care committee at the Hospital for Sick Children and were conducted in accordance with the Canadian Council on Animal Care guidelines.

### Touchscreen Training

Food restriction commenced one week prior to the beginning of behavioural training. Four days prior to the start of the experiment, all mice were handled daily for 5 min by an experimenter.

Under food restriction, each mouse received approximately 3 g of Purina rodent chow per day following their training sessions, and weights were monitored daily to ensure that no mouse dropped below 90% of *ad libitum* weight. Food restriction was in effect for the duration of the experiment. Mice were pre-trained in the touchscreen followed by training on the PAL task prior to manipulation of neurogenesis and subsequent retention testing and reversal training (Figure 1A)

**Figure 1.**
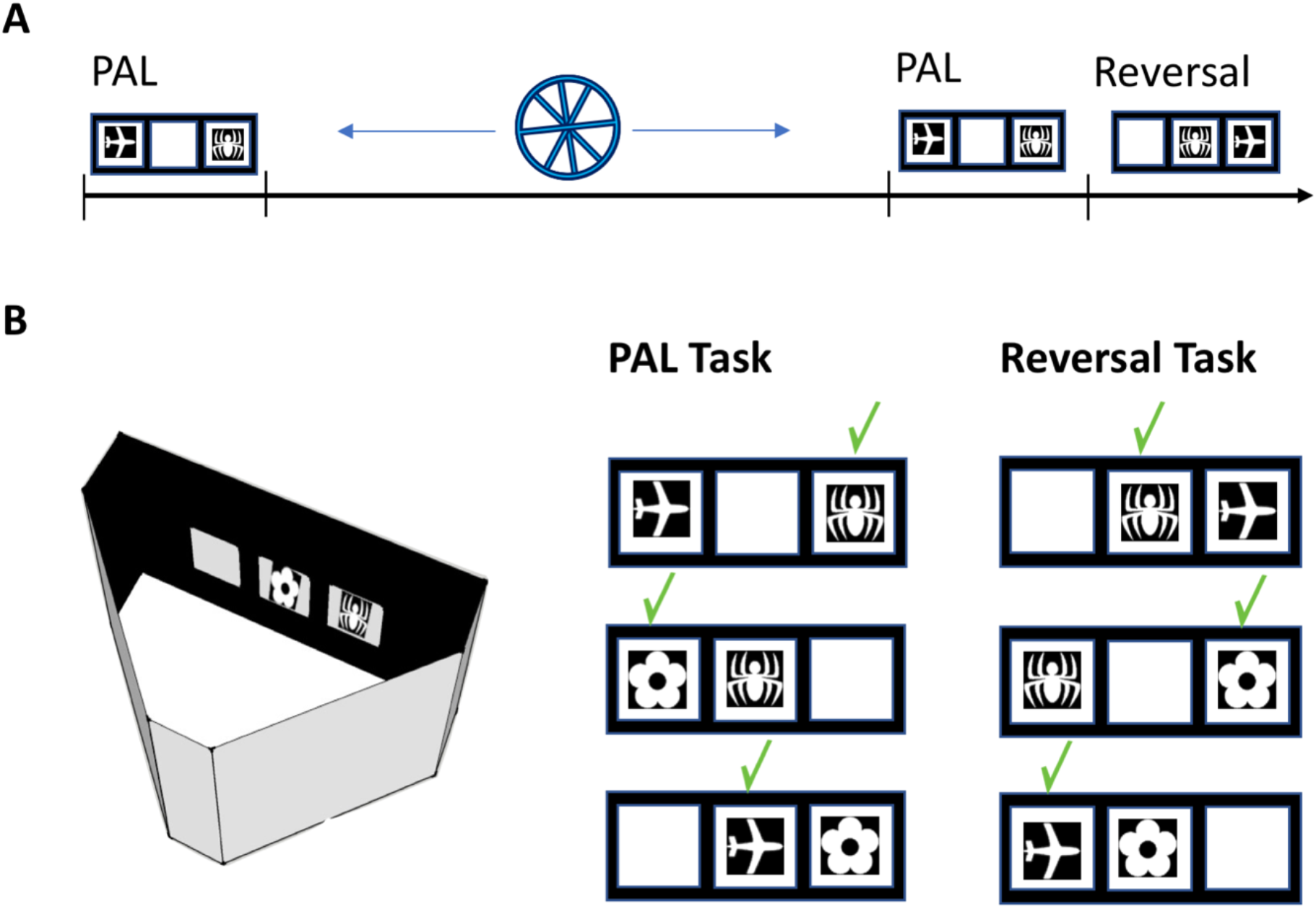
A) Experimental timeline. Mice were trained on the paired associates task untl a criterion of 80% correct was achieved. The mice were then given voluntary access to a running wheel or remained sedentary for 4 weeks. Then the mice were tested for retention of the PAL task for 7 sessions followed by reversal training in the PAL task. B) The touchscreen apparatus and the correction location of each image during the initial PAL training as well as the reversal condition.

#### Apparatus

Four touchscreen-equipped automated operant chambers for mice (Bussey-Saksida Touch Systems, Lafayette Instrument, Lafayette, Indiana) were used (Figure 1). Each apparatus consisted of a triangular modular operant chamber housed within a sound- and light-attenuating box (40 × 34 × 42 cm). The modular operant chamber consisted of a metal frame, black Plexiglas walls, and a stainless-steel grid floor. The box was fitted with a fan (for ventilation and masking of extraneous noise) and a peristaltic pump-based liquid reward well, which was illuminated by a 3-W light bulb and fitted with a photocell head-entry detector. A 3-W house light and tone generator were fitted to the back wall of the chamber.

At the front of the box opposite the liquid reward well was a flat-screen monitor equipped with an infrared touchscreen (16 cm high and 21.20 cm wide; Craft Data Limited, Chesham, UK) running ELO touchscreen software (ELO Touchsystems Inc.). Given that the touchscreen used infrared photocells, the mouse was not required to exert significant pressure on the screen for a response to be detected. A black Plexiglas slide-in mask with response windows was placed over the screen through which the mouse could make a response. The mask (h 11.80 cm × w 22.8 cm) had three response windows (h 5.8 cm × w 5.8 cm) which were positioned 0.8 cm apart from one another and located 3.0 cm from the sides of the mask.

#### Habituation

Mice were exposed to the operant chamber for 15 min per day for 2 days during which the touchscreen was blank and the three-zone response mask was in place.

#### Light-Reward training

Seven days following the start of food restriction, mice began light-reward training. Each session commenced with the delivery of 25 µl of strawberry milkshake (Neilson Dairy, Toronto, Ontario) into the reward well concomitant with the illumination of the reward well light and the presentation of a 3Khz tone of 1 sec duration. After a head entry into the reward well was detected, the light was extinguished and a fixed-interval (10 sec) schedule of reinforcement was initiated during which illumination of the reward well and sounding of the tone indicated the availability of 7 µl of strawberry milkshake. Mice received one 15 min session per day for 3 days during which the touchscreen was blank and the three-hole response mask was in place.

A decrease in Reward Latency (latency of reward well head entries following illumination of the reward well light) and an increase in the number of rewards obtained within a session reflected learning of the light-reward association.

#### Touch-Light-Reward training

Mice were trained to touch an achromatic visual stimulus presented on the touchscreen one at a time in one of the three locations of the three-hole response mask. During each trial, the location of the presented stimulus was pseudorandomized and remained on the screen until a touch to the stimulus was made. A touch response to the stimulus resulted in delivery of 7µl of strawberry milkshake in the reward well, illumination of the reward well light, and presentation of the concomitant 3Khz tone of 1 sec duration. There was a 5 sec inter-trial interval (ITI) which commenced as soon as a head entry into the reward well was detected. Responses to a blank location had no effect. The stimuli used were from a bank of 40 achromatic computer-generated visual stimuli (5 cm x 5 cm) created by the Bussey-Saksida Touch Systems (Lafayette Instrument, Lafayette, Indiana). To facilitate learning, a small amount of confectioner’s sugar moistened with water was applied to the touchscreens (in the center of the three response windows). Each daily session consisted of 30 trials or a maximum of 30 min, whichever came first. Training continued until all mice within a cage reached a performance criterion of 30 trials within a 30 min session on non-consecutive days. A further decrease in Reward Latency, a decrease in Touch Latency (latency to touch a stimulus once it appeared on the touchscreen), and an increase in the number of trials completed within a session reflected learning of the touch-light-reward association.

#### Paired-Associate Learning (PAL) task training

The PAL task required mice to learn three stimulus-place associations (Figure 1B). The three-hole response mask was employed such that there were three possible locations, each one associated with one of three visual stimuli (5 cm x 5 cm). The flower image was correct in the left position, the airplane in the central position and the spider in the right position (Figure 1B). Each PAL task training session commenced with the delivery of 7 µl of strawberry milkshake (Neilson Dairy, Toronto, Ontario) into the reward well concomitant with the illumination of the reward well light and the presentation of a 3 Khz tone of 1 sec duration. After a head entry into the reward well was detected, the light was extinguished and following a 5 sec ITI, the first PAL trial began with presentation of two of the three visual stimuli, one in the correct location and the other in an incorrect location. The third location remained blank and responses to it had no effect. If the correct stimulus was touched, both stimuli disappeared and a delivery of milkshake (7 µl) in the reward well occurred concomitant with illumination of the reward well light and presentation of the 3 kHz tone of 1 sec duration. There was a 5 sec ITI which commenced as soon as a head entry into the reward well was detected. If the incorrect stimulus was touched, both stimuli disappeared and the house light was illuminated for the duration of a 5 sec timeout period, which was then followed by a 5 sec ITI. Each session consisted of 36 trials with a maximum of 30 min. There were six different trial types with each one occurring six times per session in a pseudorandomized order such no more than three trials of the same type occurred consecutively. A correction trial occurred when an incorrect response was made and consisted of presentation of the exact same trial until a correct response occurred. Performance on correction trials did not count towards behavioral performance measures. Behavioural measures for PAL task performance consisted of percentage accuracy and number of correction trials. PAL training continued until all mice within a cage reached a performance criterion of 80% accuracy with all 36 trials completed per session. It took 50 daily training sessions for all mice to reach this level of performance.

#### Paired-Associate Learning (PAL) task reversal training

The reversal task was conducted in the same way as the PAL task itself except that the correct locations of the stimuli were changed as shown in Figure 1B. The airplane was correct when presented in the left position, the spider in the middle position and the flower in the right position. We conducted reversal training over 40 daily sessions.

### Manipulation of adult neurogenesis

To increase neurogenesis we provided half of the mice with free continuous access to a running wheel (Med-Associates) in their home cages (n=16) for a period of 4 weeks. Upon first introduction to the running wheel, each mouse was gently placed on the wheel as habituation to ensure the mice would use the running wheels. To monitor running activity, the wheels were connected wirelessly to a computer, which recorded the number of revolutions per cage. The other half of the mice (n=15) remained sedentary in standard mouse cages.

#### Histology

Mice were deeply anaesthetized with 4% chloral hydrate and were perfused transcardially, first with 40 ml of 0.1 M PBS followed by 40 ml of 4% buffered formaldehyde. The brains were then removed from the skull and post-fixed in the same 4% formaldehyde solution for 24 hours at 4°C. The fixed brains were then cryo-protected using 30% sucrose in 0.1 M PB for at least 72 hours. The tissue was rapidly frozen and sectioned in a 1/6 series at a thickness of 50 µm on a Leica cryostat. Free floating sections throughout the entire rostral-caudal extent of the hippocampus were collected and stored at -20 °C in an antifreeze solution containing 30% ethylene glycol and 20% glycerol in 0.1M PB until immunohistochemistry was performed. To quantify increases in neurogenesis one series of tissue was labeled for doublecortin. Sections were incubated in goat anti-doublecortin diluted 1:200 in 0.3% triton-x and 4% normal donkey serum for 48 hours. The tissue was rinsed 5 times for 10 minute each in 0.1 M PBS. Then, the sections were incubated for 24 hours in a 1:500 dilution of donkey antigoat Alexa488 in 0.1 M PBS, followed by 5 washes in PBS. The sections were counterstained with DAPI, mounted on glass slides and coverslipped with PVA-DABCO mounting medium. Doublecortin-labeled cells were counted using an epi-fluorescent microscope (Nikon Eclipse) and 40x objective. Cells were quantified throughout the entire dentate gyrus by an experimenter blind to treatment condition.

#### Statistical Analysis

Data was analyzed using the software package, Statistica v13.3 (Tibco). Doublecortin labeling and change in accuracy were analyzed using unpaired t-tests. Accuracy and number of correction trials for retention and reversal were analyzed using repeated measures ANOVA with Newman-Keuls post hoc tests applied for multiple comparison testing. Linear regression analysis and graphing of data were performed in Prism v8 (GraphPad).

## Results

### Acquisition of the paired associates touch screen task

To determine the impact of post training manipulations of neurogenesis on a visual paired associates task we first trained mice to achieve a criterion of 80% correct. All mice reached the 80% criterion by the end of training. Acquisition of the task occurred at a gradual rate over the course of 50 training sessions (Figure 2A). During this time, there was a corresponding decrease in the number of correction trials that were performed during each training session (Figure 2B). At the completion of training, half of the mice were given running wheels while the other half were housed in standard sedentary conditions. A significant regression equation was identified for the accuracy (F(1,47) = 519.4 *p* < 0.0001 R^2^ = 0.92) and number of correction trials (F(1,47) = 341.5 *p* < 0.0001 R^2^ = 0.88).

**Figure 2.**
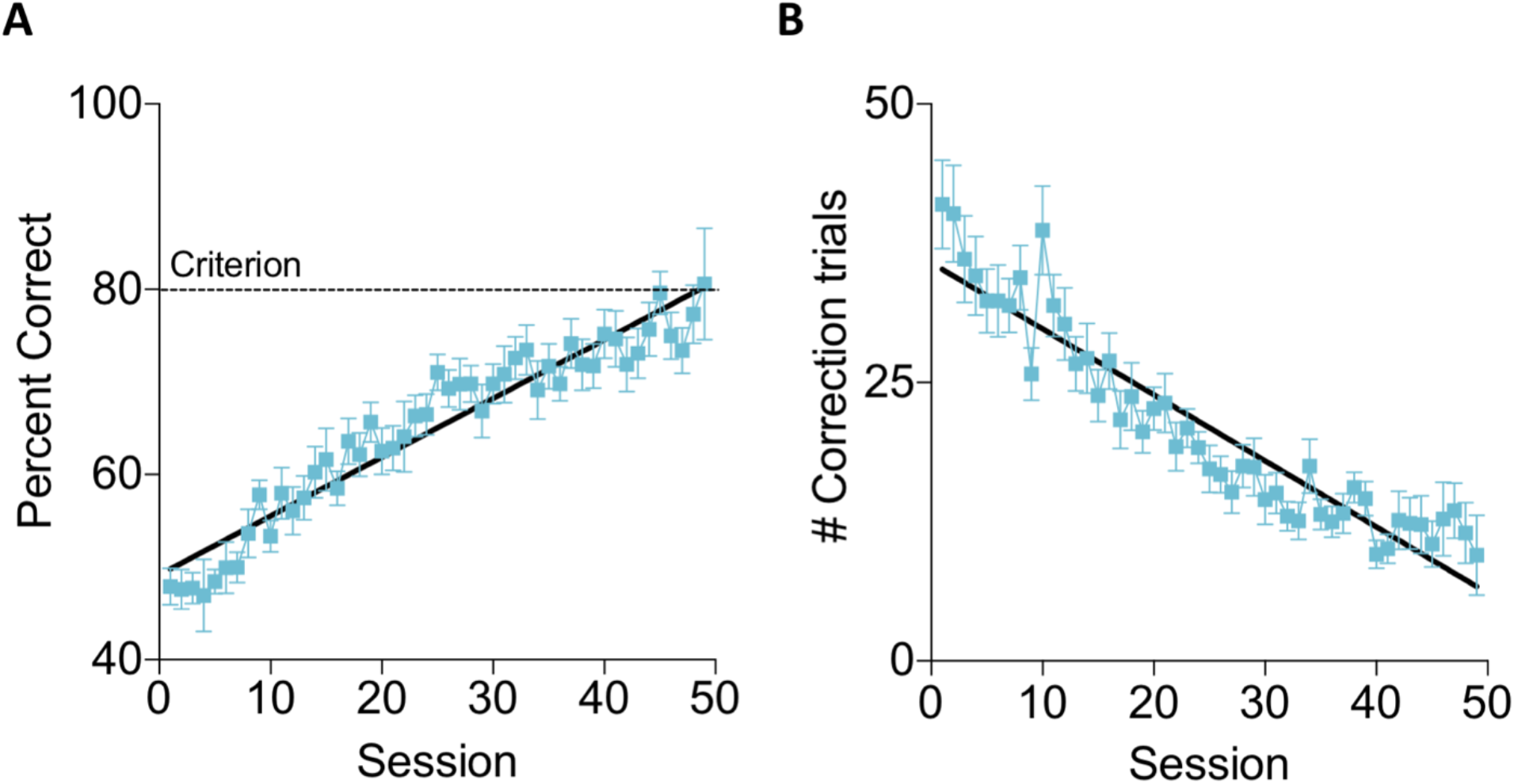
Acquisition of the PAL task. A) the accuracy of mice increased over the course of 50 training sessions and B) the number of correction trials required during each session decreased over the course of training. Data shown are mean ± SEM

### Mice that had access to running wheels showed increased forgetting of the PAL task

We reintroduced the mice to the paired associates task to assess memory retention. In both the sedentary and runner groups there was an initial drop in performance following the 4 week break from training. However, as shown in Figure 3A, the decrease in performance was significantly greater in the runner group compared to the sedentary group (t(29) = 2.03, *p* = 0.05). Our results indicate that mice with enhanced neurogenesis due to voluntary running showed impaired retention of the previously acquired PAL task (Figure 3B). There was a significant main effect of session (F(6,174) = 11.21, *p ≤ 0*.*000001*) during the 7 days of PAL retention testing indicating that performance improved over the week. There was also a significant main effect of group during the PAL retention testing sessions (F(1,29)=24.20, *p=*0.000032) indicating that the runners performed significantly better than the sedentary group. Furthermore, mice in the runner group performed significantly more correction trials during the PAL retraining sessions than did the sedentary mice (Figure 3C; F(1,29) = 22.89, *p =* 0.000046). There were no significant session by group interaction effects for either the accuracy or correction trial measures (*p’s ≥ 0*.*22)*

**Figure 3.**
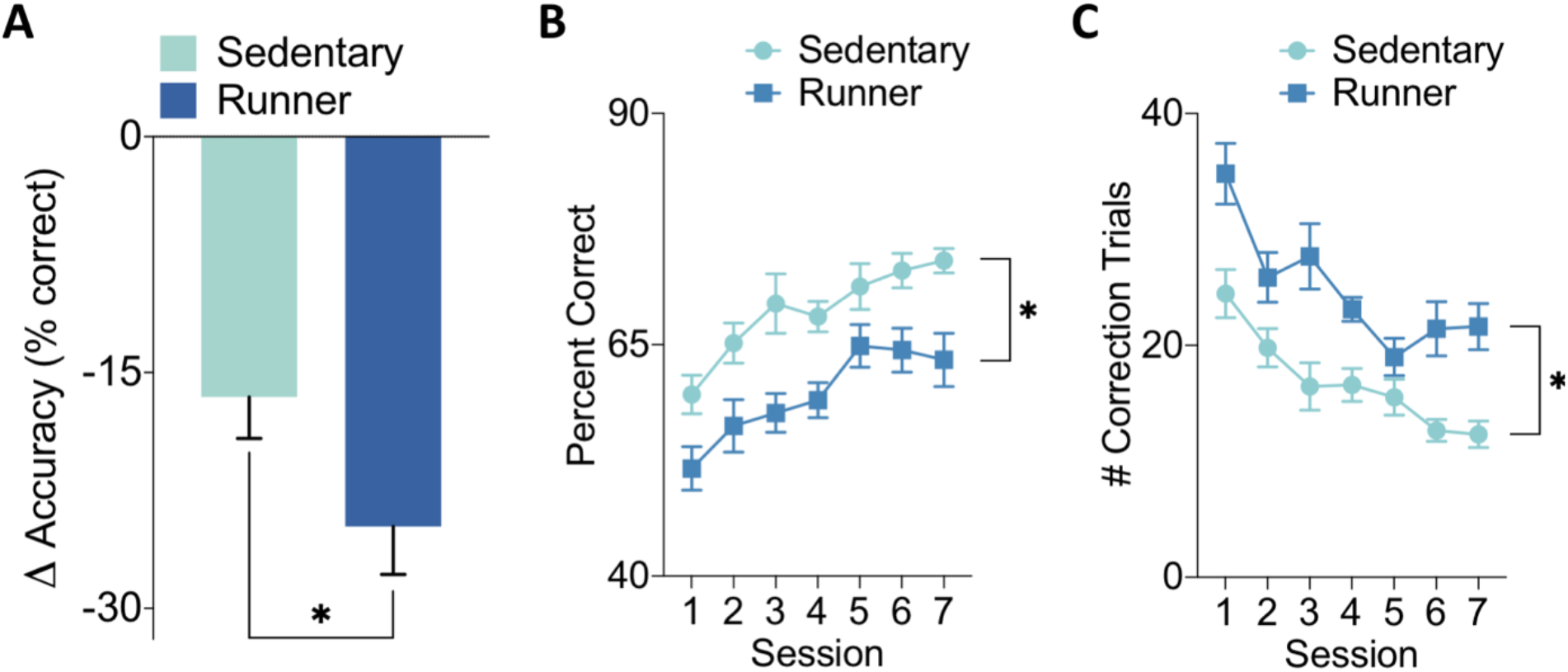
Post-manipulation PAL retention. A) The change in accuracy from the final Pre-manipulation sessions to the first post-manipulation session was significantly greater in runners than sedentary mice. B)Sedentary mice performed significantly better than runners during the 7 days of post-running testing. C)Runners used significantly more correction trials during the 7 days of PAL testing. Data shown are mean ± SEM

### Post-training manipulation of neurogenesis does not alter reversal accuracy

Following 7 days of PAL re-training we trained the mice on a PAL reversal task. The acquisition of the reversal task was not significantly different between runner and sedentary groups based on the percent correct each session (Figure 4A). There was no significant main effect of group (F(1,22) = 0.009, *p = 0*.*93)* or session by group interaction (F(39,858) = 0.90, *p = 0*.*65* but there was a significant main effect of session (F(39,858) = 18.87, *p ≤ 0*.*000001)* indicating that both groups improved over time on the reversal task Compared to the initial rate of learning there was no significant difference in the slope of the acquisition (F(1,76) = 1.59, *p* = 0.21) but the elevation of the regression line was significantly different between the acquisition and reversal conditions (F(1,77) = 59.83, *p* <0.0001).

**Figure 4.**
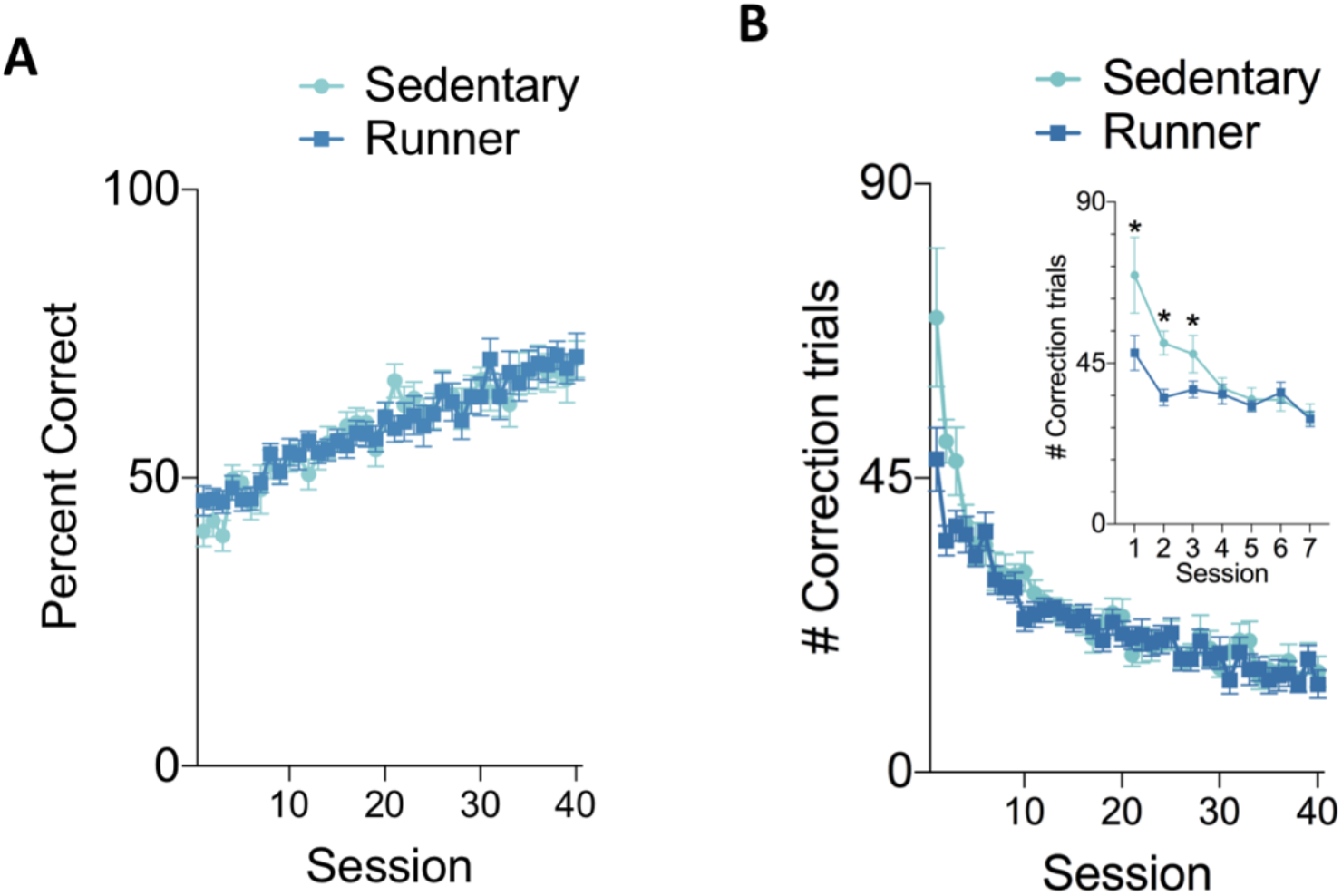
PAL reversal training. A) Accuracy during reversal learning was not significantly different between runner and sedentary mice B) However, runners required significantly fewer correction trials during reversal training. The difference was significant during the first 3 training sessions as shown in the inset figure. Data shown are mean ± SEM

### Runners performed fewer correction trials during reversal training

We assessed whether sedentary and runners differed in the number of correction trials that were required during the reversal training phase because this could indicate differences in perseveration that are dependent on the residual strength of the previous memory (Figure 4B). The number of required correction trials decreased with training (main effect of session: F(39,858) = 26.53, *p≤*0.000001). We observed that sedentary mice used significantly more correction trials during the reversal task (session by group interaction: F(39,858) = 1.63, *p* = 0.0099). The difference between groups was significant during the first three sessions of reversal training (post-hoc tests p ≤ 0.03).

### Voluntary running increases adult neurogenesis

During the 4-week neurogenesis manipulation period mice that had access to running wheels ran an average of 4.5±0.2 km per day (Figure 5A). In a separate group of age-matched mice (n=10 sedentary, n=10 runner) we used doublecortin immunohistochemistry as a measure of immature neurons to confirm that running increased neurogenesis (Figure 5B). Mice that had voluntary access to running wheels had significantly more new neurons in the dentate gyrus than sedentary mice (t(8) =2.88,*p=*0.01). This analysis was performed in a separate group of mice because the running intervention was stopped after 4 weeks and our behavioural measurements continued for another 7 weeks. At this remote time-point the initial increase in neurogenesis would have returned to baseline

**Figure 5.**
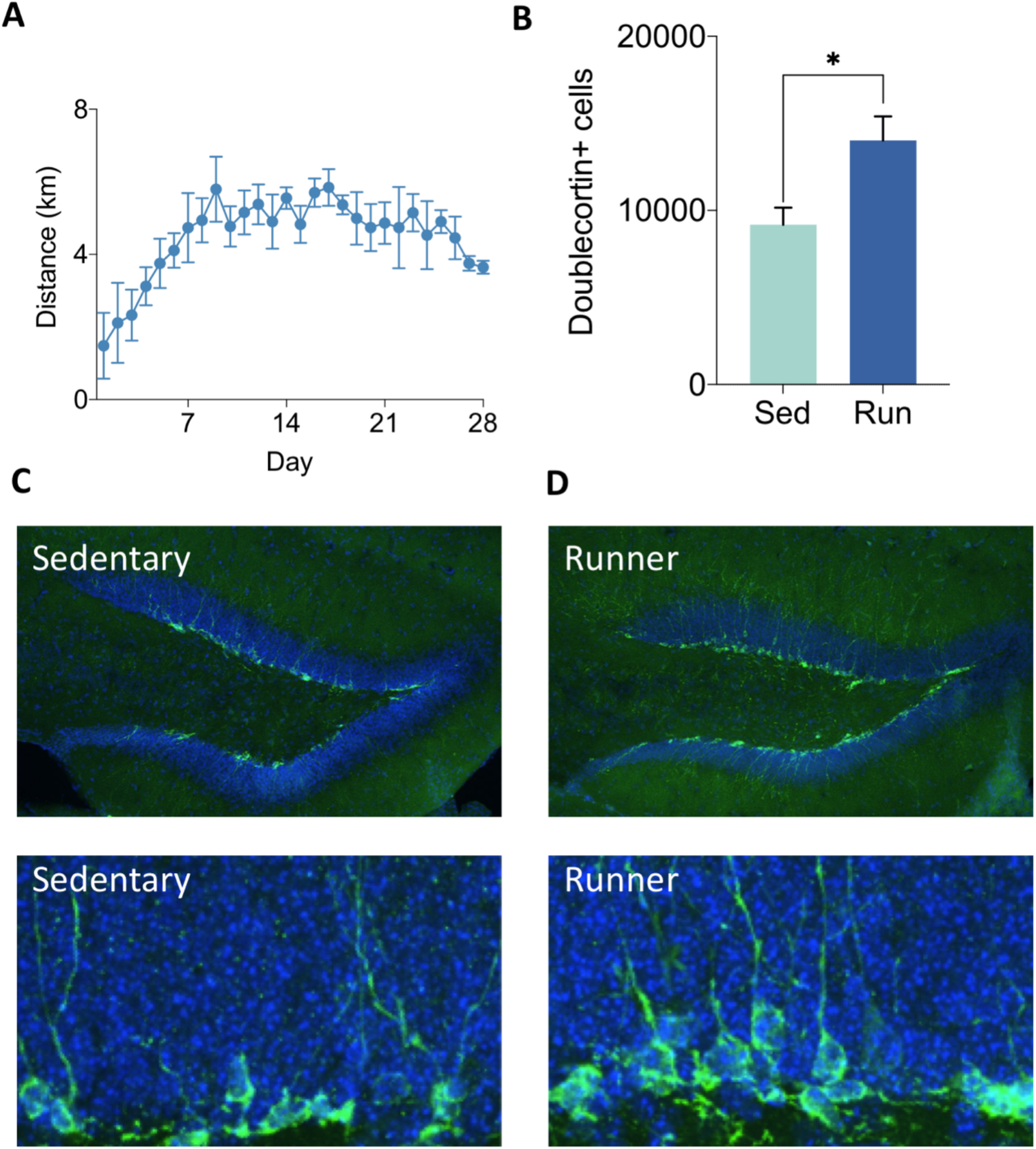
**A)** Average distance run per mouse per day during the 4 week running wheel exposure. B) Voluntary running led to a significant increase in the number of doublecortin labeled cells (immature neurons) compared to sedentary mice. C) Representative low and high magnification doublecortin labeling in Sedentary and D) runner mice. Data shown are mean ± SEM

## Discussion

Our results provide support for previous findings that elevated levels of neurogenesis induce forgetting of a previously acquired memory. Specifically, we show here that in a touchscreen based paired associates task mice with enhanced neurogenesis showed poor retention of the previous memory. We also observed some evidence in support of a reduction in interference between old and new memories as measured here by a reduction in the number of correction trials required during reversal learning. Interestingly, although we observed a significant decrease in the number of correction trials used during the initial stages of the reversal task by mice with increased neurogenesis, we did not observe a facilitation in the accuracy of this task. This is contrary to what we have previously reported on other hippocampal dependent tasks. We speculate that the difference in the current study relates to the higher degree of difficulty involved in the PAL reversal task.

In this study, we used voluntary exercise to increase neurogenesis. This manipulation results in a large enhancement in the number of new neurons produced in the dentate gyrus. Importantly, There are also other possible changes that occur in response to exercise including upregulation of growth factors and increased blood flow. While beyond the scope of the current study, we have previously determined that the forgetting which occurs because of post-training voluntary running is specifically due to the enhancement in neurogenesis (Epp et al., 2016; Scott et al., 2021). In those previous studies, pharmacological or transgenic approaches were applied to block the increase in neurogenesis while still providing running wheel access to dissociate between effects of neurogenesis or other effects of running. If neurogenesis was not elevated, wheel running did not induce forgetting. Therefore, the most likely factor that induced forgetting in the current experiment was the running indiced increase in neurogenesis.

Some previous studies have demonstrated that pre-training increases in neurogenesis facilitate different aspects of learning and memory. Enhanced memory formation has been shown to occur through several potential mechanisms including enhanced pattern separation (Clelland et al., 2009; Luu et al., 2012; Winocur et al., 2012) and increased ability to discriminate between similar memory traces (Swan et al., 2014) as well as by improving cognitive flexibility(Garthe et al., 2009, 2016; Burghardt et al., 2012). It is important to note that our findings of neurogenesis mediated forgetting are not inconsistent with this literature. In fact, in our previous study we also demonstrated enhanced learning in the Morris water task in animals that had elevated levels of neurogenesis before training.

The critical factor appears to be when the increase in neurogenesis takes place relative to the learning episode. Pre-training increases in neurogenesis may facilitate learning while post-training increases in neurogenesis promote forgetting. Post-training neurogenesis has been observed to increase forgetting in a variety of species including mouse rat, degu and guinea pig and numerous tasks such as water maze, contextual fear conditioning, Barnes maze, inhibitory avoidance and odor-context learning(Akers et al., 2014; Epp et al., 2016; Ishikawa et al., 2016; Gao et al., 2018; Cuartero et al., 2019; Scott et al., 2021) But, see also (Kodali et al., 2016).

Likely, this is due to the integration of the new neurons with the existing hippocampal circuitry; an idea which is supported by computational models(Deisseroth et al., 2004; Weisz and Argibay, 2012). If previously established patterns of connectivity exist in the hippocampus that are important for memory retrieval then the integration of new neurons would likely be disruptive but may provide additional plasticity to facilitate new encoding.

Forgetting often carries a negative connotation but it should be viewed as an important and healthy aspect of learning and memory. In fact, the importance of forgetting has been previously acknowledged. Memory storage is costly in terms of resources but many memories, while initially important become unnecessary over time. In some instances, these outdated memories can even interfere with the ability to store and retrieve updated or new memories. Therefore, inherent mechanisms that result in forgetting should not be viewed as problematic for learning and memory but instead as critical regulators of future plasticity. Our previous work has demonstrated the importance of neurogenesis induced forgetting for mitigating proactive interference (Epp et al., 2016). In that study the forgetting of previous memories allowed for facilitated acquisition of new conflicting information. In the current study we also see evidence for enhanced encoding flexibility, in terms of a reduction in the number of correction trials used during reversal learning.

Memory requires circuit stability, reductions of which may result in memory interference or poor recall. Conversely, learning requires plasticity to efficiently encode new experiences. It is clear that both processes occur and that brain regions such as the hippocampus play an important role in both learning and memory. It is less clear how the different requirements for learning and memory are regulated within the brain. Our current findings add to a growing body of research that suggests adult neurogenesis may be viewed as a mechanism to balance the demands of learning versus memory by allowing for efficient initial encoding followed by reduction in memory strength over time such that older memories may be forgotten.

## Author contributions

J.R.E., L.B.., S.A.J. and P.W.F conceived and designed the experiments. J.R.E. and L.B. performed the experiments. J.R.E conducted the analyses. J.R.E., L.B., S.A.J. and P.W.F wrote the paper.

## Acknowledgements

The authors thank Anna Gianlorenco for technical assistance with behavioral experiments. This work was supported by grants from the Canadian Institutes of Health Research to P.W.F. (FDN143227) and S.A.J. (MOP74650). J.R.E. received support from the Hospital for Sick Children and the Brain & Behavior Research Foundation.

